# Inferring cell dynamics in stress-induced neuroblastoma cell cultures

**DOI:** 10.1101/2025.03.17.643601

**Authors:** Khrystyna Hafiichuk, Sara Hamis, Jenny Karlsson, David Gisselsson, Subhayan Chattopadhyay

## Abstract

Neuroblastoma is characterised by significant intratumoural heterogeneity which complicates treatments. Phenotypic plasticity, i.e., the ability of cells to alter their phenotype without genetic mutations, is a key factor contributing to this heterogeneity. In this study, we quantify cell actions and cell-to-cell interactions that lead to phenotypic adaptation in vitro under stress, specifically low-nutrient conditions. We initially record dynamic cell counts under various nutrient conditions for two neuroblastoma cell lines with known phenotype differences. Subsequently, we compile a list of plausible biological processes that could transpire within the aforementioned systems. We then construct candidate models employing mass-action kinetics. To quantitatively infer processes that lead to phenotypic adaptation, we perform a model selection based on computational Bayesian inference methods.Our results suggest that cell-to-cell interactions promote phenotypic adaptation and that the rate of this adaptation increases with decreasing nutrient concentrations. We perform flow cytometry-based experiments to corroborate changes in phenotype composition in response to both nutrient-deprived and treatment-induced stress conditions.

## Introduction

Neuroblastoma (NB), a malignancy of the sympathetic nervous system, is one of the leading causes of pediatric cancer mortality.^1^ Although more children are now surviving neuroblastoma,^2^ it still exhibits a wide spectrum of clinical outcomes, ranging mostly from clinically responsive to aggressive, high-risk therapy-resistant metastatic disease and rarely spontaneous regression.^3^ Often resulting in poor prognosis, global treatment resistance is accepted to be the primary force behind most NB related death. Although pangenetic mutations, particularly dysregulation of the Ras / MAPK signaling pathway, have been shown to be associated with therapeutic resistance, emerging evidence suggests that phenotypic plasticity plays a crucial role in allowing the niche of NB to adapt to environmental stress.^4^

NB cells in immortalized lineages have been shown to have three morphologically distinct phenotypes capable of interconversion: neuroblastic (N), substrate-adherent (S), and intermediate (I) cell types.^5^ The S-type cells exhibit slow proliferation *in vitro*, linked to high apoptosis rates (occurring within 2–3 hours), and typically die after a few passages. They also show poor anchorage-independent growth. In contrast, N-type cells are more aggressive, proliferate robustly *in vitro*, and survive beyond 10 passages.^6^ In contrast, the N-type cells are more tumorigenic than the S-type.^7^ Recently, researchers have identified two distinct neural stem cell (NSC) states through epigenetic profiling of promoter-enhancer interactions and single-cell RNA sequencing. These states facilitate temporary adaptation to stimuli and are reminiscent of the N- and S-type differentiation states (adrenergic state: ADRN; mesenchymal state: MES), which also exhibit distinct growth characteristics.^8^ While ADRN cells demonstrate accelerated proliferative properties and exhibit a higher response to conventional chemotherapies (S-like), MES cells exhibit a dedifferentiated primitive scencent-like phenotype that is resistant to treatments (N-like).

While the majority of high-risk neuroblastoma tumors initially exhibit an ADRN phenotype making them vulnerable to targeted treatment such as anti-GD2 antibody therapy, a portion of tumors present with mesenchymal predominance.^9,10^ Furthermore, investigation into the nature of the two states show that NB cells are able to switch between these states based on the tumor environment, a process referred to as *phenotypic plasticity*.^11^ The fluid interconversion between ADRN and MES states is very likely responsible to some extent for the emergence of treatment resistance in neuroblastoma. This phenotypic plasticity of ADRN and MES states results in temporal variations in their admixture in response to dynamic stress conditions, which subsequently influences the population growth characteristics of NB cells.

We hypothesize that by observing these growth characteristics in NB model systems of varied phenotypic makeup, it may be possible to estimate the relative proportions of cells in the two states, provided that state conversion rates are realistically accounted for. Previous studies in insulinomas demonstrate that estimating the relative proportions of two phenotypes interacting in a closed system, based on growth dynamics, can reveal treatment resistance patterns *in vitro*.^12^ Our research aims to extend beyond the point estimates from a snapshot to uncover the underlying mechanistic rules governing phenotypic plasticity of NB model system. In doing so, we seek to theoretically predict the growth characteristics of a system based on initial observations under generalized stress conditions.

## Results

### Model selection suggests that cell-to-cell interactions promote phenotypic adaptation in low-nutrient neuroblastoma cell cultures

To study neuroblastoma cells under stress, we cultured cells from the SK-N-BE(2)C and IMR-32 cell lines in various low concentrations of fetal bovine serum (FBS). Cell count data from these experiments demonstrated non-monotonic growth curves in which cell counts initially increased during a growth phase, reached a peak, and subsequently decreased (**Figure 1a-b**). To quantitatively infer the cell actions and cell-to-cell interactions that gave rise to these growth curves, we performed a mathematical model selection based on fitting candidate ordinary differential equation (ODE) models to data via a Hamiltonian Monte Carlo Sampling (HMCS) algorithm implemented in the probabilistic programming framework Stan (see Methods for details). The selected model that best captured the data is pictorially illustrated in **Figure 1c**. The model follows a two-phenotype convention in which the cells are in one of two plastic phenotype states: state-1 or state-2. Respectively, these describe a more and a less proliferative state and could be likened to either N-type and S-type cells, or to noradrenergic and mesenchymal cells. Our results suggest that cell-to-cell interactions catalyse state-1 to state-2 cell conversions that are key in driving the non-monotonic growth curve dynamics (**Figure 1c**). Moreover, our simulation results show that both peak heights and peak time occurrences are correlated with nutrient concentrations, such that they generally increase with increasing FBS concentrations (**Figure 1d**).

**Figure 1:**
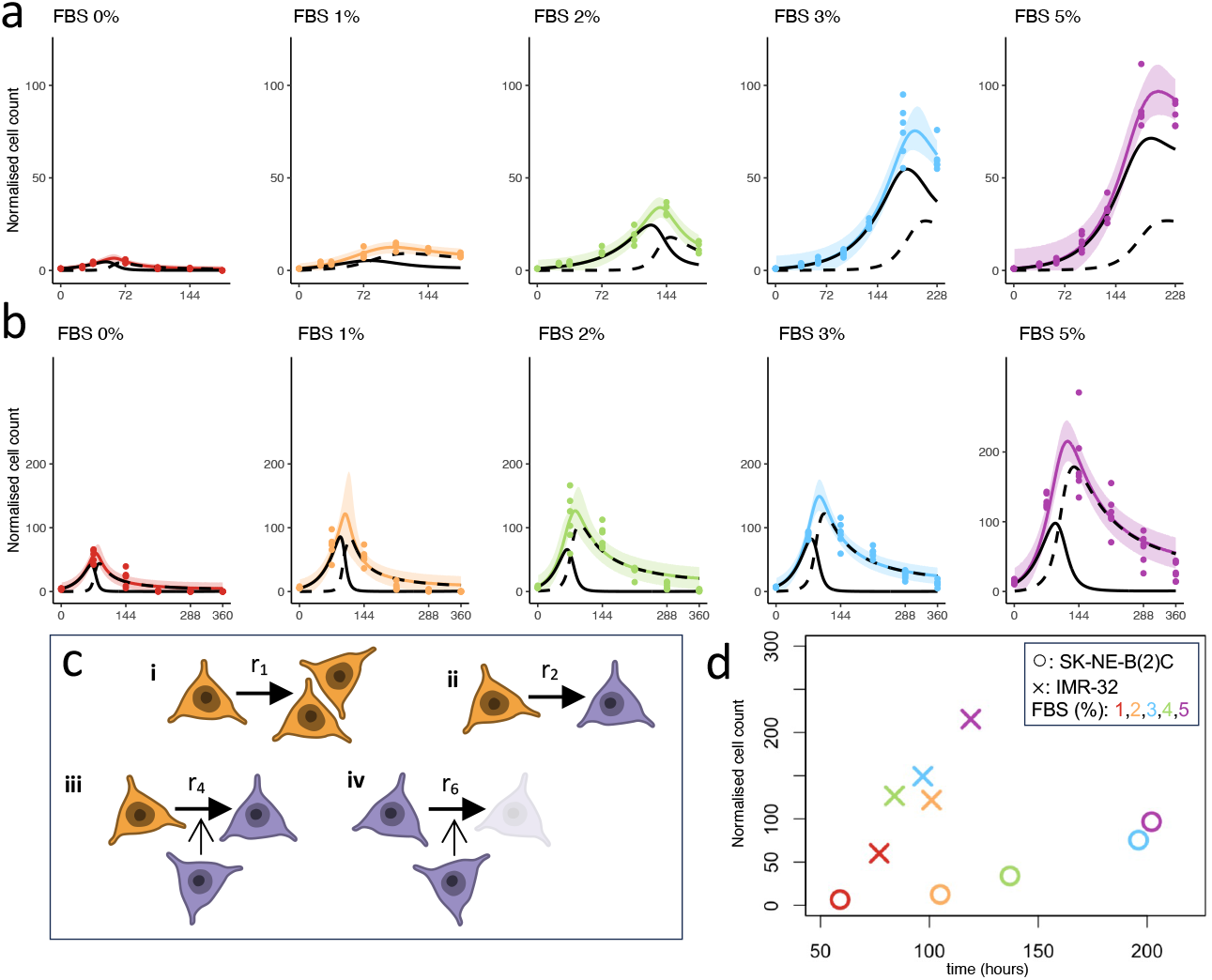
Data matched simulations suggest that cell-to-cell interactions impact dynamics in neuroblastoma cell cultures. In vitro cell counts (points) and modelled cell count dynamics (means and 90% credible intervals in coloured lines and bands, respectively) are shown for (a) SK-N-BE(2)C and (b) IMR-32 cell cultures. Solid and dashed black lines show modelled state-1 and state-2 counts, respectively, normalised at 0 hours. (c) The inferred model processes describe actions and interactions of cells in state-1 (orange) and state-2 (purple). The processes are: (i) state-1 cell division, (ii) spontaneous state 1-to-2 cell state conversion, (iii) state-2 induced conversion from state-1 to state-2 and (iv) state-2 induced death of cells in state-2. (d) Modelled peak heights are plotted against modelled peak times.

### Model selection suggests that decreasing nutrient concentration speed up cell plasticity and death in neuroblastoma cell cultures

To visualise how the rates of the biological processes in the selected model (**Figure 1c**) depend on FBS concentrations, we plot posterior probability density distributions of the inferred model parameters (Figure 2). These distributions do not show a strong correlation between the rate of cell division (*r*_1_) and FBS concentrations between 0 and 5%. However, the distributions demonstrate that *r*_4_, the rate of phenotypic adaptation induced by cell-to-cell interactions, decreases with increasing FBS concentrations for both SK-N-BE(2)C and IMR-32 cells. These results suggest a nutrient dependence of cell-to-cell interactions in neuroblastoma which, in turn, impact cell plasticity and survival. Furthermore, in IMR-32 but not SK-N-BE(2)C cells, we see a positive correlation between spontaneous cell state conversion (*r*_2_) and FBS concentrations, and a negative correlation between interaction-induced cell death (*r*_6_) and FBS concentrations. Also showin in Figure 2b are the inferred posterior probability density distributions of the normalised initial count of cells in state-1 (*y*_1,IC_) and state-2 (*y*_2,IC_), which were not observable in the in vitro experiments.

**Figure 2:**
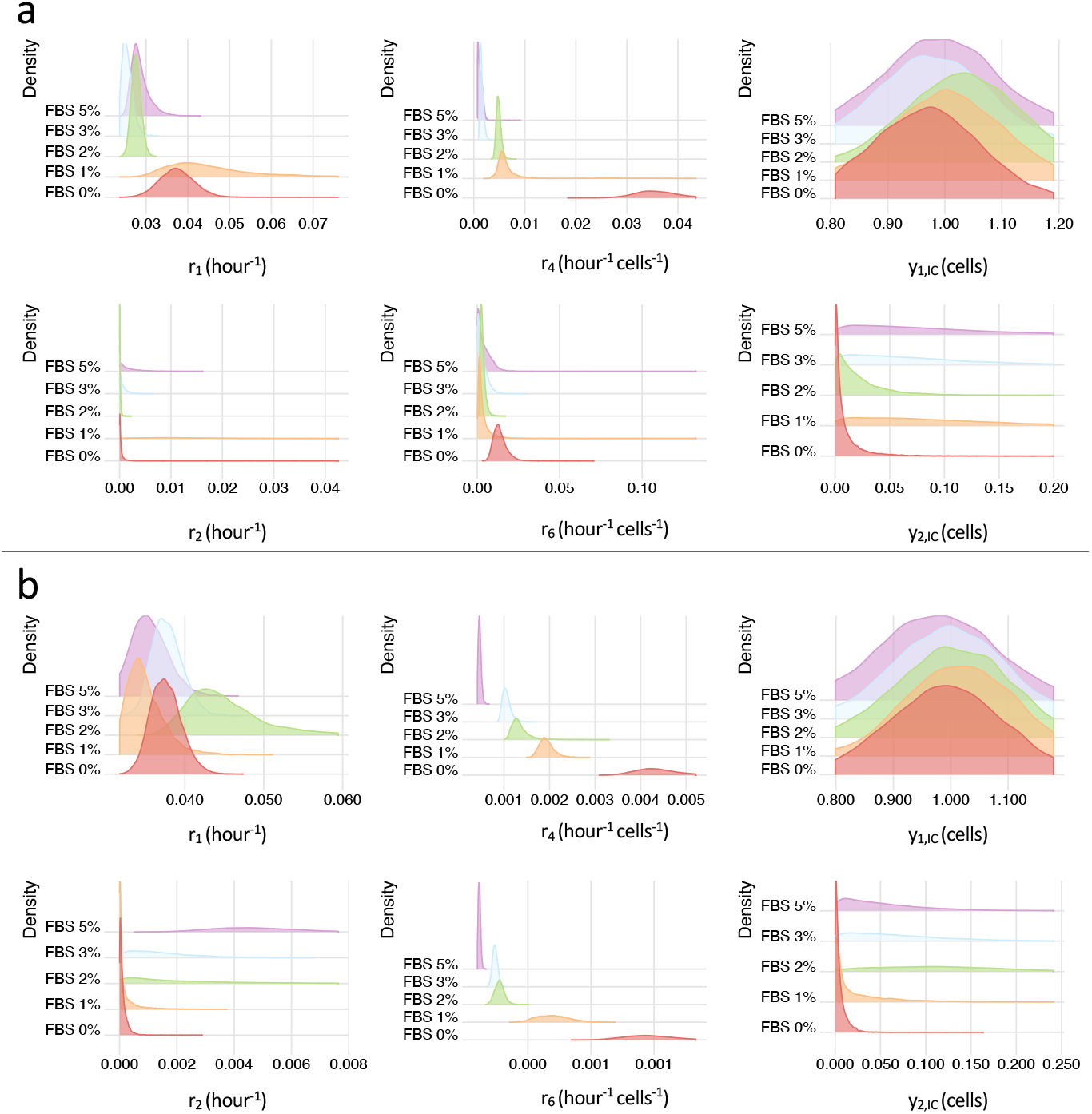
Parameter inference suggests that the rate of phenotypic adaptation increases with decreasing nutrient concentrations. Posterior probability densities are shown for model parameters in (a) SK-BE-N(2)C and (b) IMR-32 cell lines. These describe spontaneous state-1 cell division (*r*_1_), spontaneous state-1-to-2 conversion (*r*2), state-2 induced conversion from state-1 to state-2 (*r*_4_) and state-2 induced death of cells in state-2 (*r*_6_). The inferred, normalised initial cell counts for state-1 (*y*_1,IC_) and state-2 (*y*_2,IC_) cells are also shown. The prior probability densities are *r*_*j*_ ∼ Half-Normal(0, 0.15^2^)∀*j* for SK-BE-N(2)C, *r*_*j*_ ∼ Half-Normal(0, 0.05^2^)∀*j* for IMR-32, and *y*_1,*IC*_ ∼ Normal(1, 0.1^2^), *y*_0,*IC*_ ∼ Half-Normal(0, 0.1^2^), with all model parameters enforced to be positive.

### Phenotypic plasticity and stress adaptation in neuroblastoma cell-lines

As we have established a relationship between environmental stress and phenotypic plasticity, we then examine how neuroblastoma cells with distinct baseline compositions respond to different stress conditions. We hypothesized that the phenotype transition represents a fundamental adaptive response to diverse stressors (see e.g, **Figure 1**), and that the initial phenotypic composition of the population critically determines the outcome of this adaptation. Previous work from our colleagues has established that SK-N-BE(2)C cells adapt a more mesenchymal (S2 like) phenotype under chemotherapy.^11^ To further test this, we subjected IMR-32 cells, which exhibit a predominantly adrenergic (S1) phenotype under wildtype conditions, to both nutrient stress as well as chemotherapeutic stress while monitoring the phenotypic changes through surface marker expression.

We use flow cytometry for characterizing neuroblastoma phenotype switching between the S1 like ADRN and S2 like MES states by enabling high-throughput, single cell analysis of surface marker expressions of CD44 and GD2 proteins. CD44, a cell surface glycoprotein involved in cell adhesion and migration, is upregulated in MES like neuroblastoma cells and is associated with therapy resistance and tumor-initiating properties. Conversely, GD2, a disialoganglioside highly expressed on neuroectodermal tumors, is a hallmark of ADRN type cells and is commonly targeted in high-risk neuroblastoma with immunotherapy. By using fluorochrome-conjugated antibodies against CD44 and GD2, we can delineate ADRN (GD2high/CD44low) and MES (GD2low/CD44high) populations and quantify shifts in phenotype.

To determine whether this phenotypic plasticity was specific to genotoxic stress or represented a more general adaptive mechanism, we repeated the experiment under nutrient deprivation conditions. Culturing IMR-32 cells in media containing only 2% FBS produced a pronounced shift toward the MES phenotype over 72 hrs (P<0.05, relative population frequency difference, Figure 3), confirming that the ADRN to MES transition also operates as a conserved survival strategy for IMR-32. Next we test effect of genotoxicity with treatment of 5 µM cisplatin for 72 hours. Here IMR-32 cells demonstrated a more rapid shift toward a mesenchymal phenotype, as evidenced by a greater than 5,000-fold increase in CD44 expression within 48 hours (P<0.001, Figure 3).

**Figure 3:**
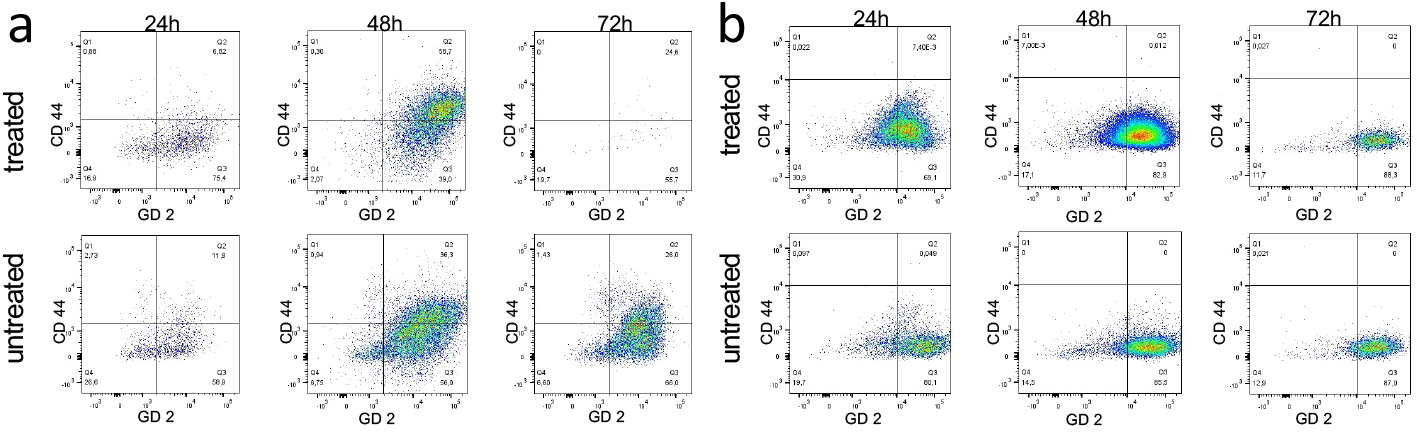
Environmental stressors affect IMR-32 cultures. (a) 5 μM cisplatin treatment over 72 hours in treated and untreated groups show relative cell populations in Q1 (CD44highGD2low), Q2 (CD44highGD2high), Q3 (CD44lowGD2high) and Q4 (CD44lowGD2low). (b) Cells in treatment group are cultured in 2% FBS media for 72 hours and surface expression of CD44 and GD2 are evaluated in a similar manner to that in (a). Both untreated groups are cell cultured in optimal growth conditions.

## Discussion

Although the standard of care chemotherapeutic agents and targeted therapies initially achieve complete response in the majority of NB, these treatments frequently become ineffective over time.^13^ This phenomenon necessitates the escalation of treatment regimens especially for high-risk groups to progressively aggressive and highly toxic combination therapies, such as rapid-COJEC, which incorporates five chemotherapeutic agents.^14^ The discovery that NB often comprises two distinct phenotypes, one of which may mediate global cross-resistance, has significantly influenced the field’s understanding of clinical outcomes and has informed the design and analysis of our preclinical experiments.

To understand the dynamic shift between these two clinically significant phenotypes, it is imperative to understand the ecological and evolutionary dynamics of cellular populations exhibiting the ADRN and MES tendencies, particularly under environmental stressors, which all therapeutic conditions can be considered surrogates of. Despite the absence of a clear consensus on the classification of these two phenotypes, the terms ADRN and MES are frequently used inter-changeably to describe both cell types and cell states. A critical distinction between these concepts is that cell types are heritable, whereas cell states are not. Recent research by our colleagues has demonstrated that SK-N-BE2(C) cells *in vitro* are prone to phenotype interconversion, with cells reverting to their original state upon removal of the stressor.^11^ This study demonstrates that in the two NB cell line model systems, the pre-exposure composition of ADRN and MES phenotypes differs post-exposure significantly. Previous investigations focusing on N and S type cells have reported differences in relative abundance of these types in IMR and BE cells. Consistent with these previously published data, we demonstrate that pre-exposure cell populations of the two cell lines comprise markedly different relative proportions of ADRN/MES, as determined by surface marker expression.

The mathematical framework presented herein demonstrates that a bi-phenotypic system proxied with cell states effectively recapitulates the heterogeneous growth kinetics observed under environmental stress in different neuroblastoma lines. In addition, the simulations reveal that phenotypic adaptation is governed by interaction-dependent state transition rates. The model also predicts divergent evolutionary trajectories based on starting conditions i.e., populations with pre-existing phenotypic heterogeneity demonstrate adaptive plasticity through stress-induced phenotype enrichment.

These theoretical predictions find direct experimental validation in our flow cytometric analyses (Figure 3). The IMR-32 cell line, characterized by ADRN dominant baseline composition, demonstrates limited adaptive capacity under both chemotherapeutic and nutrient stress, ultimately progressing to apoptotic elimination which stands in contrast to the more phenotypically diverse SK-N-BE(2)C line, which is shown to maintain population viability through dynamic MES expansion (quantified by CD44 upregulation). Yet, IMR-32’s shift toward mesenchymal phenotype under different stress conditions were statistically significant compared to untreated controls and accompanied by a corresponding decrease in GD2 expression, mirroring observations reported in SK-N-BE(2)C cells. However, the absence of a substantial pre-existing MES subpopulation in IMR-32 cells (approximately 10% MES in wildtype) appeared to limit their adaptive capacity, as evidenced by widespread apoptosis by 72 hours of treatment. This contrasts with the behavior of phenotypically heterogeneous SK-N-BE(2)C (with approximately 40% MES in wildtype), which can maintain viable MES populations under prolonged stress due to their intrinsic cellular diversity. Thus, we note that the attenuated response under nutrient stress compared to chemotherapy may reflect differences in the intensity or nature of the selective pressures involved. Importantly, the eventual collapse of the IMR-32 population under both stress conditions highlights the critical role of initial phenotypic diversity in determining adaptive outcomes.

These findings collectively establish a mechanistic framework wherein neuroblastoma cell populations navigate environmental stress through interaction-driven phenotypic remodeling, with population survival contingent upon both the starting phenotypic distribution and the kinetics of interconversion between cellular states.

The experimental component of this study is confined to *in vitro* experiments using only two cell lines which we acknowledge needs expansion to elucidate a more generalized conclusion. In addition, as the initial conditions of the ADRN/MES make up affected the results, the inflection point of population rescue is a joint function of the initial condition and environmental stress levels which remain unexplored. While our findings are consistent with previously published data and shed light on mechanistic relationship of phenotype switch, further investigations are needed to demonstrate clinical generalizability of these results and how ecological stressors are translated in treatment contexts. As the predominance of either of the phenotypes would imply a tumor following recourse in accordance to the response rates if unchanged from standard of therapy, despite the novel mechanisms of ecological herding discussed in this text, evaluating the phenotype make up would add great clinical value to NB prognostication.

## Methods

### Cell cultures and cell counting

SK-N-BE(2)C and IMR-32 cells well cultured in FBS concentrations of 0%, 1%, 2%, 3%, 5%. For SK-N-BE(2)C cells, the cell viability of four to six samples was measured at seven time points between 0 and 228 hours for each FBS concentration (**Figure 1a,b**). For IMR-32 cells, the cell viability of six samples was measured at six time points between 0 and 360 hours for each FBS concentration (**Figure 1b**).

Cell viability was assessed using the CountessTM Automated Cell Counter from Invitrogen with Trypan blue exclusion. Cells were mixed with 0.4% Trypan blue solution (1:1 dilution), and live cells were quantified using a TC20 Automated Cell Counter (Bio-Rad). For each sample, counts were performed in duplicate. To obtain total cell counts, cell viability was multiplied by culture volume. After each sampling, FBS concentrations were replenished in the cell cultures.

### Flow cytometry

For flow cytometric analysis, 50,000 cells were seeded per well in a 6 well plate. After 24 hours, cells were either treated with 5 μM cisplatin (prepared in PBS), 2% FBS, or maintained as untreated controls. Following treatment, cells were harvested by trypsinization, washed in PBS, and stained with fluorophore-conjugated antibodies in 100 μL FACS buffer (PBS, 0.5% BSA, 4 mM EDTA) for 30 min in the dark. After staining, cells were washed (PBS followed by FACS buffer) and resuspended in FACS buffer containing DAPI (1:1000 dilution) for viability discrimination. Samples were acquired on a BD LSR Fortessa flow cytometer. Each experiment included compensation controls, fluorescence-minus-one (FMO) controls, and isotype controls. Technical triplicates were analyzed for all conditions and estimates were calculated pooling all replicates.

### Formulation of mathematical candidate models

In our model, we abstractly classify cells as being in one of two phenotype states, state-1 or state-2, where the former state is more aggressive than the latter. We denote the densities of cells in these states at time *t* by *q*_1_(*t*) and *q*_2_(*t*), respectively. The states can be related to the classical neuroblastoma N-type and S-type cells, or be partially aligned with the more recently categorised noradrenergic and mesenchymal cells. As a first step towards inferring which underlying cell actions and cell-to-cell interactions may give rise to the experimental cell population behavior observed in **Figure 1a-b**, we formulate a list of theoretically motivated permissible biological processes that might occur in the cell cultures (**Table 2**). As spatial structures are perturbed in the cell cultures after sampling, and we do not have spatial data, we make the simplifying modelling assumption that all cells in a vial are equally likely to interact with each other. With this assumption, we can use the law of mass action to formulate each permissible biological processes as an ODE. From the list of permissible processes, we construct candidate models such that each candidate model comprises a unique set of permissible processes with at least one permissible process related to cell division, one to phenotypic adaptation, and one to cell death. Each candidate model is thus described by a system of ODEs, in which each term in the equation right-hand-sides describes the dynamics of a permissible process.

We denote each permissible processes by an index *p*, and each candidate model by an index *m*. The set *P* ^*m*^ ⊂ {1, 2, …, *N*} is unique to each candidate model *m*, and describes which of the *N* candidate processes *p* = 1, 2, …, *N* are included in the model (here *N* = 6). For each candidate model, the cell culture system is described by the state vector 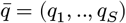, where *S* is the number of phenotypes (here *S* = 2). Thus, the *m*th candidate model is described by a system of ODEs

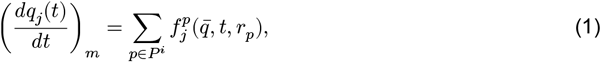

where *m* = 1, 2, …, *M* and *M* is the number of candidate models (here *M* = 8). From **Table 1**, we see that, for instance, candidate model 4 is described by four permissible processes: cell division (*p* = 1), spontaneous cell state conversion (*p* = 2), *q*_2_-induced cell state conversion (*p* = 4) and *q*_2_-induced cell death (*p* = 6). Therefore, *P* ^4^ = {1, 2, 4, 6} and the model equations are

**Table 1:** Antibody information table.

| Antibody | Conjugation | Clone | Lot Number | Manufacturer | Species | Dilution |
| --- | --- | --- | --- | --- | --- | --- |
| CD44 | AF488 | BJ18 | B320781 | Biolegend | mouse | 1:50 |
| GD2 | AF647 | 14.G2a | 562096 | BD | mouse | 1:25 |
| igG2a K | AF488 | 20102 | IC003G | R & D Systems | mouse | 1:50 |
| igG2b K | AF647 | MPC-11 | 400330 | Biolegend | mouse | 1:50 |

**Table 2:**
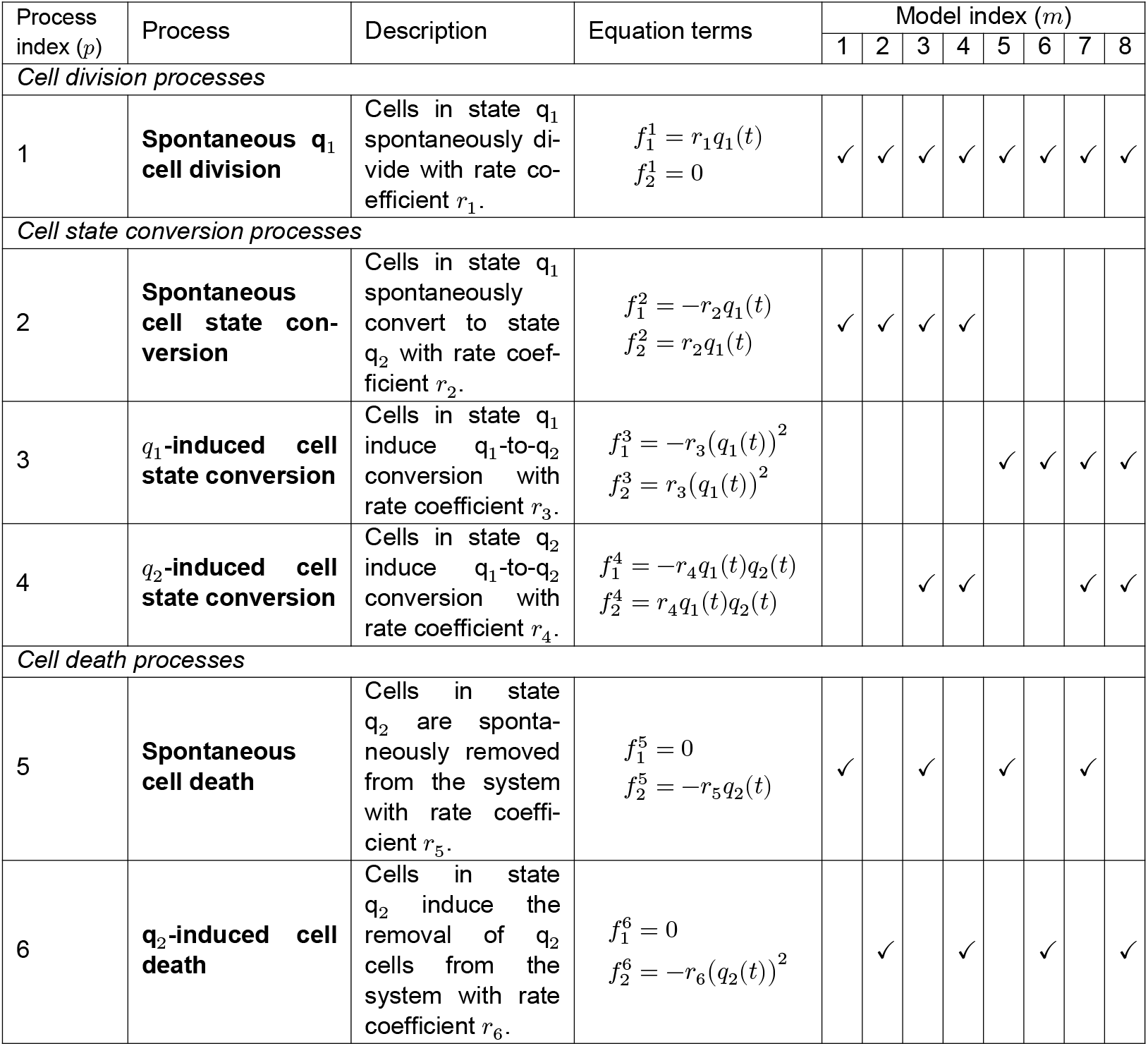
Permissible biological processes and candidate models. The table illustrates which permissible processes 1 to 6 are included in candidate models 1 to 8. Included processes are indicated by check marks (✓).

| Process index ($p$) | Process | Description | Equation terms | Model index ($m$) | | | | | | | |
| --- | --- | --- | --- | --- | --- | --- | --- | --- | --- | --- | --- |
|  |  |  |  | 1 | 2 | 3 | 4 | 5 | 6 | 7 | 8 |
| Cell division processes |  |  |  |  |  |  |  |  |  |  |  |
| 1 | Spontaneous $q_1$ cell division | Cells in state $q_1$ spontaneously divide with rate coefficient $r_1$. | $f_1^1 = r_1 q_1(t)$<br>$f_2^1 = 0$ | ✓ | ✓ | ✓ | ✓ | ✓ | ✓ | ✓ | ✓ |
| Cell state conversion processes |  |  |  |  |  |  |  |  |  |  |  |
| 2 | Spontaneous cell state conversion | Cells in state $q_1$ spontaneously convert to state $q_2$ with rate coefficient $r_2$. | $f_1^2 = -r_2 q_1(t)$<br>$f_2^2 = r_2 q_1(t)$ | ✓ | ✓ | ✓ | ✓ | | | | |
| 3 | $q_1$-induced cell state conversion | Cells in state $q_1$ induce $q_1$-to-$q_2$ conversion with rate coefficient $r_3$. | $f_1^3 = -r_3 (q_1(t))^2$<br>$f_2^3 = r_3 (q_1(t))^2$ | | | | | ✓ | ✓ | ✓ | ✓ |
| 4 | $q_2$-induced cell state conversion | Cells in state $q_2$ induce $q_1$-to-$q_2$ conversion with rate coefficient $r_4$. | $f_1^4 = -r_4 q_1(t) q_2(t)$<br>$f_2^4 = r_4 q_1(t) q_2(t)$ | | | ✓ | ✓ | | | ✓ | ✓ |
| Cell death processes |  |  |  |  |  |  |  |  |  |  |  |
| 5 | Spontaneous cell death | Cells in state $q_2$ are spontaneously removed from the system with rate coefficient $r_5$. | $f_1^5 = 0$<br>$f_2^5 = -r_5 q_2(t)$ | ✓ | | ✓ | | ✓ | | ✓ | |
| 6 | $q_2$-induced cell death | Cells in state $q_2$ induce the removal of $q_2$ cells from the system with rate coefficient $r_6$. | $f_1^6 = 0$<br>$f_2^6 = -r_6 (q_2(t))^2$ | | ✓ | | ✓ | | ✓ | | ✓ |

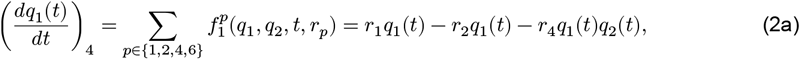

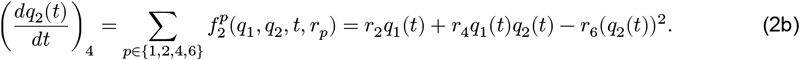

Note that, in formulating the candidate models, we have taken a minimal-parameter modelling approach, such that we have models that contain exactly one cell division and death process, and only uni-directional cell state conversion processes. This is equivalent to assuming that there is one dominating cell division and cell death process, and one main direction of phenotypic adaptation in the constant FBS concentration. Another model simplification is that the rate coefficients *r*_*k*_, *k* = 1, …, *p* implicitly may depend on FBS concentrations. However, we assume that the rate coefficients are not functions of time since (1) FBS concentrations are replenished after sampling in the experimental setup, and thus FBS concentrations are assumed to be constant in time, and (2) the cells do not run out of space in the wells, and thus we assume that no space-mediated cell-to-cell competition occurs.

### Implementation and model selection

The measured experimental observable is the total cell count over time, with no information about the composition of cells in state-1 and state-2. In order to, despite this, make progress in uncovering dynamics in the modelled cell system, we heuristically fit all candidate models to the in vitro data using the probabilistic programming framework Stan and RStan, the R interface to Stan. A visual model selection (Supplementary Material; SM1) reveals that the models that are best able to capture the observed in vitro dynamics are models 4 and 8. These include three common processes: cell division of state-1 cells (*p* = 1), state-2 induced state-1-to-2 cell state conversion (*p* = 4), and state-2 induced state-2 death (*p* = 6). Further, model 4 includes spontaneous state-1-to-2 cell state conversion (*p* = 2), while model 8 includes state-1 induced state-1-to-2 cell state conversion (*p* = 3). Since process 2 results in linear ODE terms (±*r*_2_*q*_1_(*t*)), and process 3 results quadratic ODE terms (±*r*_3_*q*_1_(*t*)^2^), model 4 is the simplest model that captures the in vitro dynamics.

