## supplementary material for "Inferring cell dynamics in stress-induced neuroblastoma cell cultures"

### Supplementary Material (SM1: Model selection)

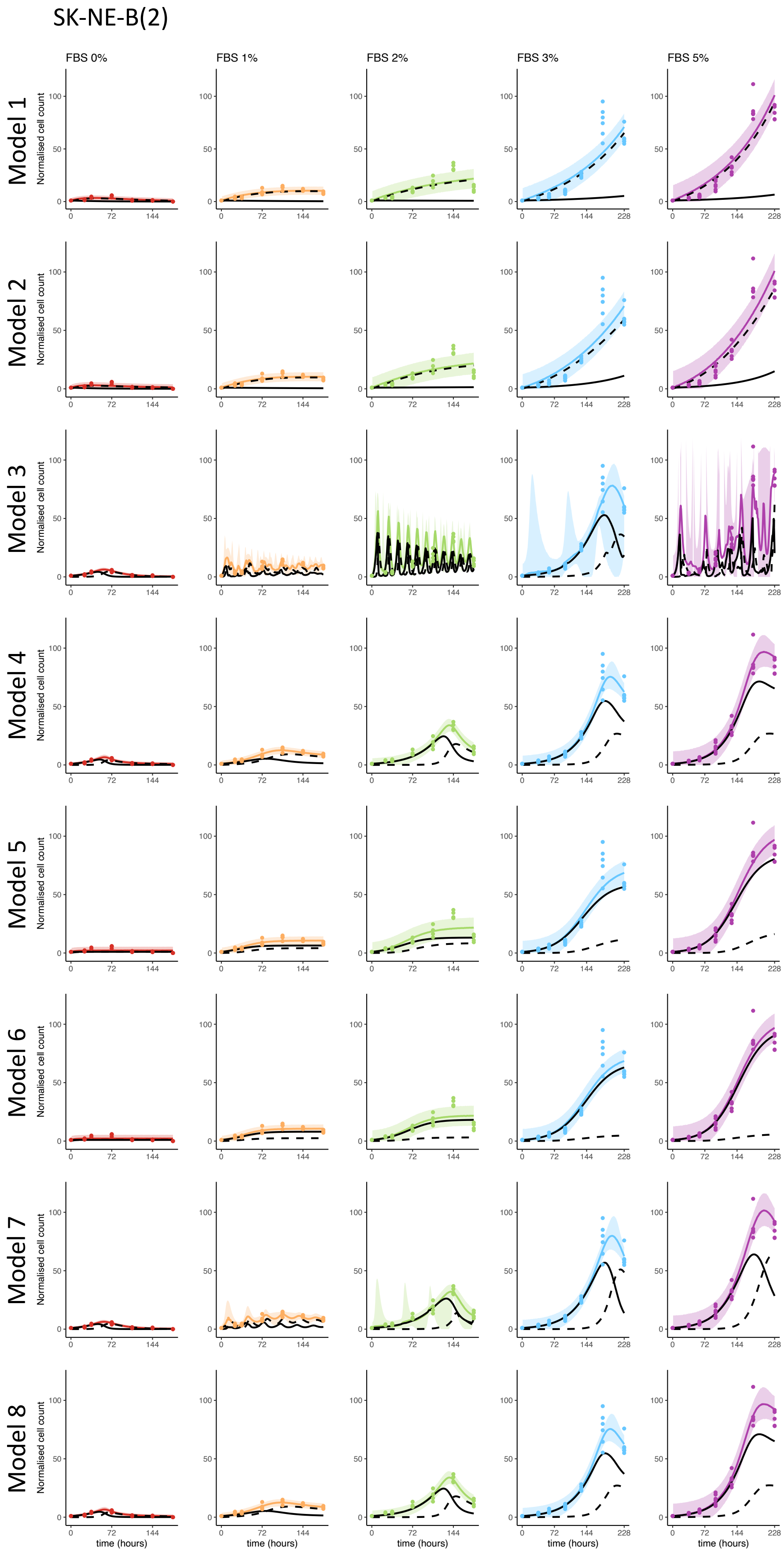


Figure 1: Model selection for SK-BE-N(2)C cells.

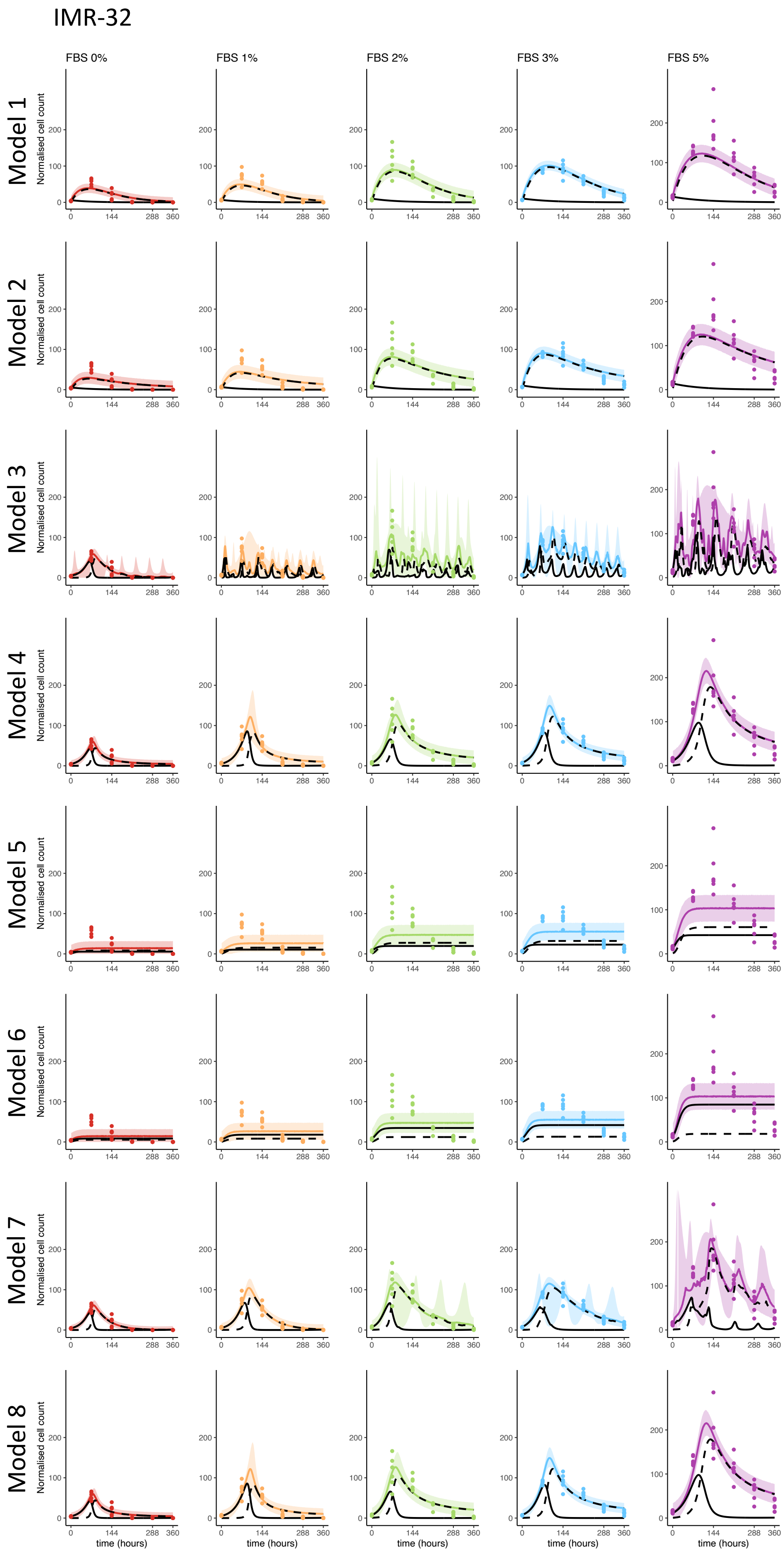


Figure 2: Model selection for IMR-32 cells.
